# High-resolution models of actin-bound myosin from EPR of a bifunctional spin label

**DOI:** 10.1101/458257

**Authors:** Benjamin P. Binder, Andrew R. Thompson, David D. Thomas

## Abstract

We have employed two complementary high-resolution electron paramagnetic resonance (EPR) techniques with a bifunctional spin label (BSL) to test and refine protein structural models based on crystal structures and cryo-EM. We demonstrate this approach by investigating the effects of nucleotide binding on the structure of myosin’s catalytic domain (CD), while myosin is in complex with actin. Unlike conventional spin labels attached to single Cys, BSL reacts with a pair of Cys; in this study, we thoroughly characterize BSL’s rigid, highly stereoselective attachment to protein α-helices, which permits accurate measurements of orientation and distance. Distance constraints were obtained from double electron-electron resonance (DEER) on myosin constructs labeled with BSL specifically at two sites. Constraints for orientation of individual helices were obtained previously from continuous-wave EPR (CW-EPR) of myosin labeled at specific sites with BSL in oriented muscle fibers. We have shown previously that CW-EPR of BSL quantifies helix orientation within actin-bound myosin; here we show that the addition of high-resolution distance constraints by DEER alleviates remaining spatial ambiguity, allowing for direct testing and refinement of atomic structural models. This approach is applicable to any orientable complex (e.g., membranes or filaments) in which site-specific di-Cys mutation is feasible.

## Introduction

Actin and myosin share a fundamental partnership within motile organisms, taking on a particularly specialized role in the function of striated muscle. Myosin couples actin-activated hydrolysis of ATP, actin binding and release, and structural changes within its own catalytic domain (CD) to produce mechanical force on actin filaments. Upon activation of muscle contraction, this biochemical and structural coupling induces relative sliding of thin and thick filaments in the myofibril lattice, shortening the fiber (1–3).

The greatest barrier to precise structural understanding of actomyosin function and pathology currently lies in difficulties associated with the analysis of both proteins in complex. No crystal structures of actin-bound myosin have been reported, and thus the resolution of atomic models is currently limited to that of electron microscopy. Moreover, both crystallography and EM are intrinsically limited to the study of virtually homogenous samples in restricted spatial environments, imposing conditions that may not accurately reflect the dynamic structural equilibrium at play during myosin’s catalytic cycle. Further refinement of existing mechanistic models thus depends on the derivation of high-resolution structural constraints using alternative methods.

Site-directed spectroscopy provides the versatility and resolution necessary for quantifying small allosteric changes within dynamic proteins. Fluorescence and electron paramagnetic resonance (EPR) have been used extensively to characterize biochemical, kinetic, and structural changes in myosin (4–7). The comparatively superior intrinsic resolution of EPR has proven especially powerful in the study of intra-myosin distances by double electron-electron resonance (DEER) (7, 8), and of the orientation of myosin domains within oriented muscle fibers (9, 10). Until recently, however, the effective resolving power of site-directed EPR has been limited by nanosecond mobility of the spectroscopic probe relative to its associated labeling site, contributing significant static and dynamic disorder to the resulting spectral waveform (11, 12).

The recent adoption of bifunctional spin labels (BSL) for site-directed labeling has shown great promise for eliminating the effect of local probe dynamics on EPR spectra (Fig. 1) (13, 14). By engineering two cysteine labeling sites at positions *i* and *i*+4 on a stable α-helix, the resulting bifunctional reaction of BSL’s twin methanethiosulfonate groups strongly anchors its paramagnetic center to the protein backbone, ensuring that the spectrum accurately reflects the behavior of the backbone (15). We have recently employed this approach in oriented muscle fibers to derive the orientations of individual myosin α-helices relative to the actin filament axis (16). The BSL approach has also been used to great advantage over traditional spin labels in the accurate determination of DEER distance distributions for multiple protein systems, both experimentally (15, 17) and *in silico* (18).

**Fig. 1.**
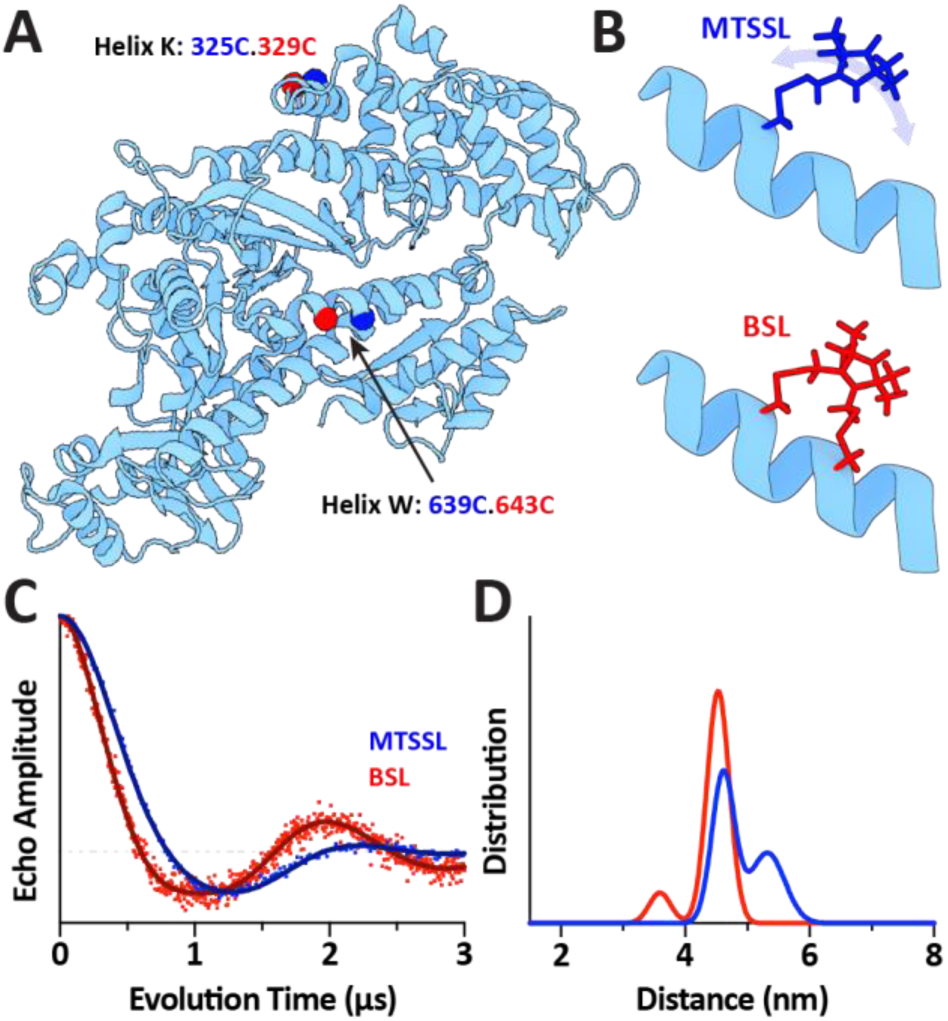
Comparison of MTSSL and BSL for distance measurements across the myosin CD. A: Structure of myosin with labeling sites highlighted. MTSSL-labeled constructs were designed by introducing cysteine residues at positions 325 (Helix K) and 639 (Helix W) (blue). BSL-labeled constructs were designed by introducing an additional cysteine residue (red) on each helix to form *i*and *i*+4 bifunctional labeling sites at positions 325.329 and 639.643. B: Illustration of binding modes for MTSSL and BSL. C: DEER waveforms for 325.639-MTSSL (blue) and 325.329.639.643-BSL (red), with simulations overlaid (dark blue and dark red). D: Best-fit Gaussian distance distributions for 325.639-MTSSL (blue) and 325.329.639.643-BSL (red) corresponding to waveforms in C.

Taken together, these high-resolution orientation and distance measurements represent a set of orthogonal structural constraints similar to those often employed in nuclear magnetic resonance for structure determination (19). Both measurements are applicable to myosin bound to oriented actin, and can be performed under a wide variety of biochemical conditions. Considered alone, neither measurement is capable of directly producing a unique atomic model for elements of the complex, but simultaneous analysis of both orientation and distance constraints has not yet been reported. In the present work, we hypothesize that the combined use of BSL in oriented fiber EPR and in DEER can significantly alleviate spatial degeneracy, and render sufficient structural information for the development of precise atomic models of actomyosin structural elements.

## Methods

### Protein preparations

Mutant myosin constructs were prepared in a “Cys-lite S1dC” *Dictyostelium* myosin II background truncated at residue 758 and containing only one native (non-reactive) Cysteine at position 655 (20). Constructs were expressed and purified from *Dictyostelium* orf+ cells. Actin was extracted from rabbit skeletal muscle acetone powder, and prepared as described previously (21).

### Site-directed spin labeling

Di-Cys and tetra-Cys myosin mutants in labeling buffer (30mM Tris, 50mM KCl, 3mM MgCl2, pH 7.5) were incubated at 4°C with 5mM DTT for 1hr to ensure reduction of engineered cysteine residues prior to labeling. DTT was removed using Zeba Spin desalting columns (Thermo Scientific), and myosin was incubated at 4°C for 1hr in 8-fold molar excess of either the bifunctional 3,4-Bis-(methanethio sulfonylmethyl) −2,2,5,5-tetramethyl-2, 5-dihydro-1H-pyrrol-1-yloxy spin label (BSL), or the monofunctional 1-Oxyl-2,2,5,5-tetramethyl-Δ3-pyrroline-3-methyl methanethiosulfonate spin label (MTSSL). Following incubation, excess spin label was removed and the protein was exchanged into EPR (rigor) buffer (40mM HEPES, 1mM EGTA, 2mM MgCl2, 15mM KPr, pH 7.0) using Zeba Spin desalting columns.

### EPR spectroscopy and analysis

DEER samples were prepared by adding 7% v/v glycerol (as a cryoprotectant) to samples containing ∼100μM spin-labeled myosin, loading samples into quartz capillaries (1.1mm i.d., 1.6 mm o.d., 15μL sample volume) and subsequently flash freezing samples in liquid nitrogen. A Bruker E580 spectrometer operating at Q-band (34 GHz) with an EN5107 resonator was then used to implement a four-pulse DEER protocol with a π/2 pulse width of 12 ns and an electron double resonance (ELDOR) pulse width of 24 ns. The ELDOR frequency was placed on the pump position, which corresponded to the absolute maximum of the nitroxide absorption spectrum. The observe position was placed 24 Gauss higher than the pump position on the field-swept absorption spectrum. Experiments were run at a temperature of 65 K.

DEER data was analyzed using custom software written in Mathematica (available for download at github.com/thompsar), integrating components of DeerAnalysis ((22), Tikhonov regularization and background validation) and DEFit ((23), Monte Carlo-based model fits). Raw, normalized waveforms were fit to a 3D-homogeneous background model and the background component was removed. Tikhonov regularization was performed on the background corrected waveform, with choice of smoothing parameter being selected by leave one out cross validation ((24), LOOCV) as well as by inspection of the L-curve. The region of background fit was then varied (i.e. background validation) to identity regions of stability in the distance distributions for background choices that yielded Tikhonov fits of comparable quality (Fig. S1, Fig. S2). To aid in further parameterization of the stable portions of the distance distribution by a Monte Carlo fit with a sum of Gaussians, regions of the distance distribution that were highly unstable based on background choice (and thus artifacts or distances beyond the sensitivity limit afforded by the evolution time), were suppressed by augmenting the background model with the necessary components to suppress the unstable peaks, with no distortion of the stable distance distribution(s) after Tikhonov Regularization of the reprocessed raw data. The distance distribution was used as a seed for a sum of Gaussians Monte Carlo fit of the background-corrected waveform, with the appropriate number of Gaussians being informed by the Bayesian Information Criterion to avoid overfitting (25). The Monte Carlo fits yielded error surface plots for the varied parameters (centers, widths and mole fractions), and confidence intervals were defined by using an f-ratio distribution for comparing the ratio of the residual sum of squares (RSS) of the entire search with the best RSS. Additional details of the analysis are discussed in Supplemental Information (Fig. S1, Fig. S2).

### Molecular modeling

Angle and distance measurements derived from atomistic models were calculated using custom extensions written for Visual Molecular Dynamics 1.9.3 (VMD) (26). Minimization of models was performed with help from the SciPy and NumPy packages written for the Python Programming Language. Structure images were rendered using Blender, Version 2.79 (Blender Foundation).

## Results

### Comparison of monofunctional and bifunctional spin labels for DEER distance measurements

Three locations on myosin’s catalytic domain (CD) were chosen for this study, at positions previously determined to give unique orientational distributions and exhibit minimal perturbation of actin-activated myosin ATPase (16). To assess potential improvement of the DEER measurement in myosin with BSL, we focused first on measuring the distance between sites on Helix K and Helix W, because these helices appear to be relatively static in myosin-only crystal structures, and give rise to narrow orientational distributions when studied individually with BSL in oriented muscle fibers (16, 27–29).

DEER waveforms of myosin containing two single-Cys motifs and labeled with MTSSL display oscillations that are significantly attenuated relative to waveforms from BSL-labeled di-Cys constructs, suggesting a more disordered population in the MTSSL-labeled samples (Fig. 1C). Waveforms from both experiments were subjected to Tikhonov fitting analysis, which was used to seed for a sum-of-Gaussians Monte Carlo fit, in order to parameterize the distance distribution (Fig. S1). The MTSSL-labeled single-Cys construct gives rise to a distance distribution best fit with two Gaussians, a primary distribution parameterized with (r1=5.16nm, σ1=0.33nm,*χ*1=0 55), and a much narrower secondary distribution parameterized with (r2=4.58nm, σ2=0.15nm, *χ*2=0.45) (Fig. 1D, blue). The BSL-labeled di-Cys construct is also best fit with two Gaussians, a primary distribution parameterized with (r1=4.58nm, σ1=0.26nm, *χ*1=0.95), and a secondary distribution parameterized with (r2=3.52nm, σ2=0.26nm, *χ*2=0.05) (Fig. 1D, blue). The two overlapping Gaussians centered at ∼4.6nm indicate that both experiments contain some fraction of labels with nearly identical relative placement, consistent with the equivalent design of the single-Cys and di-Cys labeling sites. However, in the BSL-labeled construct, the narrow 4.58nm distribution comprises 95% of the total population, while in the MTSSL-labeled construct, the equivalent 4.58nm distribution comprises only 45% of the total population. We hypothesize that the small secondary distribution in the BSL experiment represents a well-defined minor conformation of the label that was previously observed in oriented fibers (16), and further evidence for this hypothesis is presented in the following sections. The MTSSL results, on the other hand, are dominated by a much broader distribution that may represent some fraction of the large number of possible rotamer states available to a flexibly-attached probe. From these results, we thus concluded that the use of BSL represents a clear advantage for high-resolution distance detection in our myosin system.

### Distances measured across the myosin CD in the presence of actin

We generated three myosin constructs using each possible pairing of the three BSL labeling sites selected for study, and recorded DEER for each labeled construct (Fig. S1). Of the three possible constructs, two produced DEER waveforms with stable distance distributions relative to background signal correction for the evolution times observed: 325.329.639.643-BSL (HK-to-HW, employed in the previous section; Fig. 2A, left), and 492.496.639.643-BSL (Relay-to-HW; Fig. 2A, right). Thus, these two constructs were selected for further study; fitting of waveforms from myosin-only samples and analysis of derived distance distributions is shown in Fig. 2B-C (blue), with associated parameters given in Table 1. When prepared alone without nucleotides or additional proteins, fitting results for the two constructs display striking similarity: both are optimally parameterized by two Gaussians, with a very narrow primary distribution and a minor secondary distribution.

**Fig. 2.**
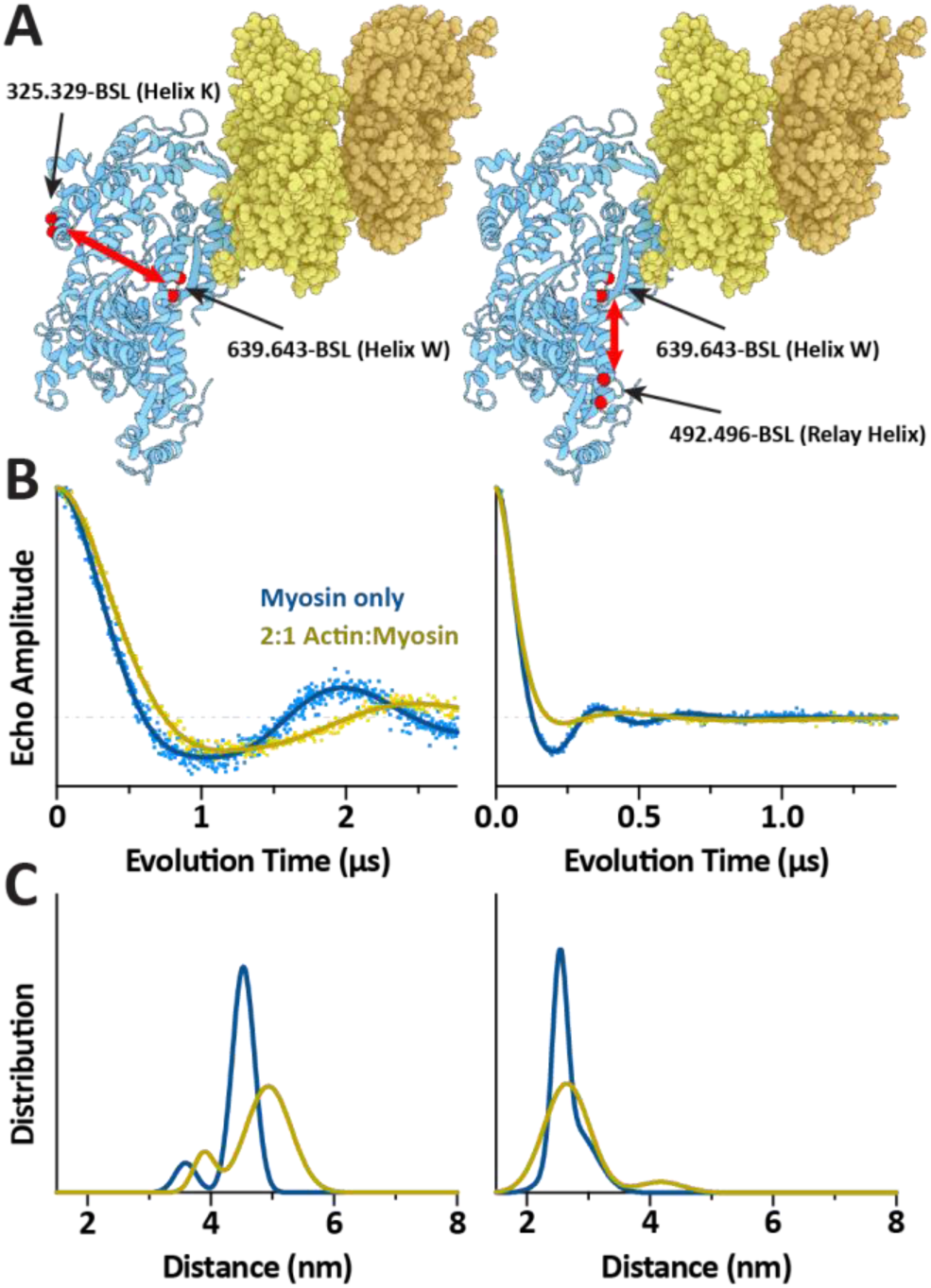
Comparison of the actin-free and actin-bound states of myosin, measured using double-BSL constructs. A: Illustrations of measured distances in the HK-to-HW construct (left) and relay-to-HW construct (right), showing two actin monomers from different chains (yellow and orange), the myosin CD (blue) and relevant BSL labeling sites (red spheres). Actin filament axis is normal to the plane of the page. B: Background-corrected DEER waveform data from the constructs illustrated in A, in the absence of actin (blue) and the presence of twofold molar excess actin (yellow). Best-fit Gaussian-based simulations are overlaid in dark blue and dark yellow, respectively. C: Gaussian distance distributions corresponding to the fits in B.

**Table 1.**
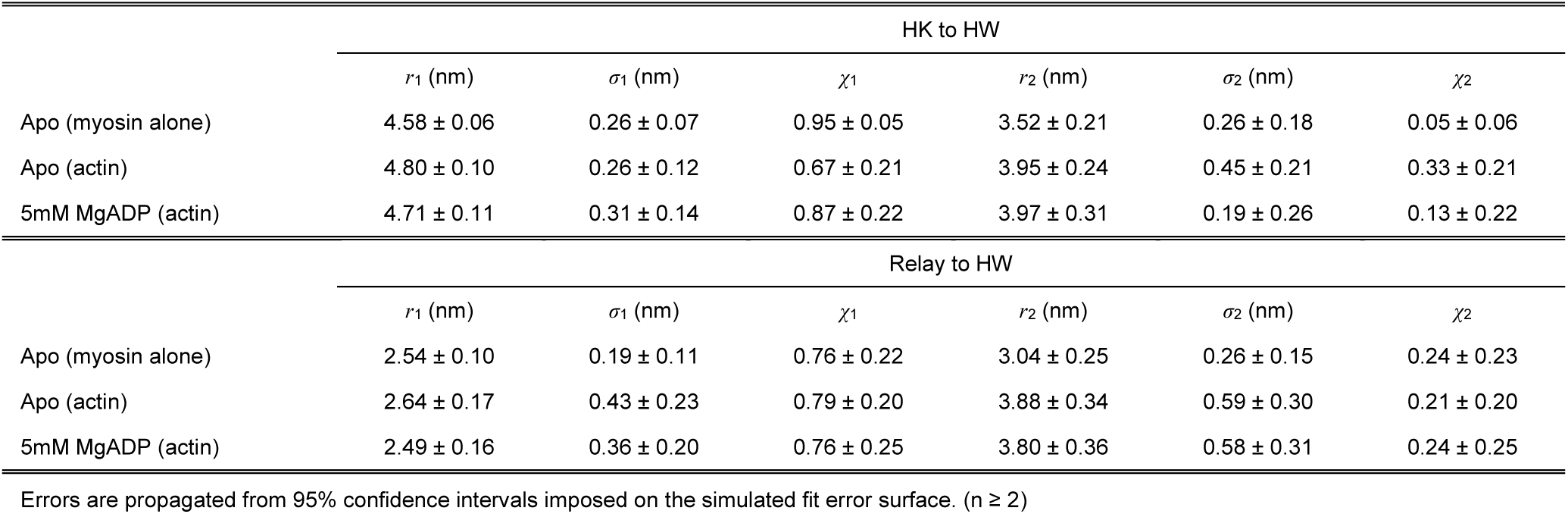
Average Gaussian parameters for DEER-detected distance distributions in actin-bound myosin.

Next, we recorded DEER on both constructs in the presence of F-actin, using a ratio of 2:1 actin:myosin (mol/mol) (Fig. 2B, yellow). Fitting analysis produced distance distributions which, as with the actin-free samples, were best parameterized by the sum of two Gaussians (Fig. 2C, yellow; Table 1). Table 1 shows that distance distributions with and without actin are generally similar, with one noteworthy difference: The HK-to-HW construct shows a significant increase in interprobe distance upon addition of actin. indicating structural changes across the actin-binding cleft. This result is in general agreement with a previous DEER study using monofunctional labels in similar locations across the cleft (8), although without additional labeling pairs in the cleft region, it is not possible to determine whether this change represents a closing or opening transition.

### No significant changes in inter-helix distance distributions are detected with MgADP

In order to assess potential changes to intra-myosin distances with nucleotide binding/release in the actomyosin complex, we recorded DEER on our relay-to-HW and HK-to-HW constructs in the presence of both actin (2:1 actin:myosin, mol/mol) and 5mM MgADP (Fig. 3B, light/dark green). Fitting analysis produced distributions that were not statistically distinct from their nucleotide-free analogs (Table 1), although the average primary distance derived from the Relay-to-HW construct does exhibit a slight negative shift relative to Apo conditions, consistent with a similar construct studied previously without actin present (7). Overall, this result is not surprising, as evidence from previous structural studies indicates that the effects of MgADP within the myosin CD probably manifest as subtle rotations of individual structural elements, with minimal relative spatial displacement (29–33). We therefore hold these inter-helix distance centers constant in the constrained modeling of the MgADP-bound rigor complex in the following section.

**Fig. 3.**
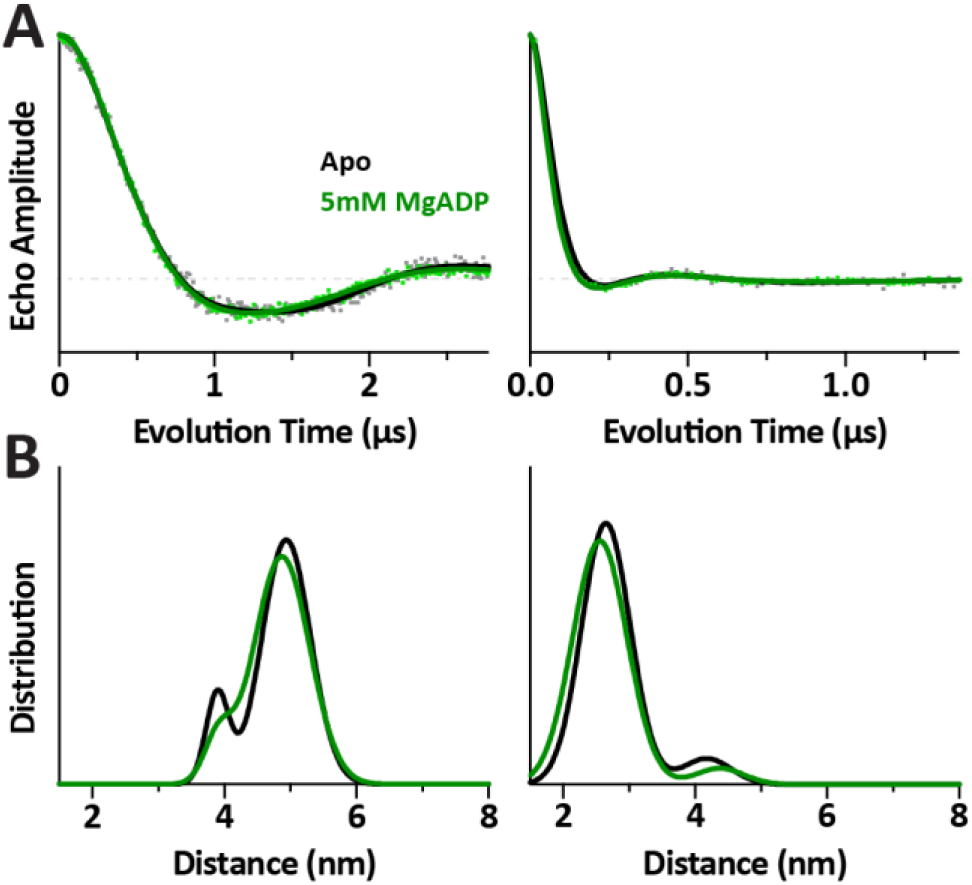
Comparison of the apo and MgADP states of the actomyosin complex, measured using two double-BSL constructs. A: Background-corrected DEER waveform data for the HK-to-HW construct (left) and Relay-to-HW construct (right), in the absence of nucleotide (gray) and the presence of 5mM MgADP (green). Best-fit Gaussian-based simulations are overlaid in black and dark green, respectively. C: Gaussian distance distributions corresponding to the fits in B.

### An atomic model of the apo complex satisfies experimental constraints, and characterizes BSL’s binding conformation

Molecular modeling was employed to compare our EPR-based measurements of inter-helix distance, as well as previous data reporting on individual helix orientations (16), with established atomistic models. Our goal was to produce a model for a BSL-labeled actomyosin complex, validated both by the literature and our experiments, which could subsequently be used as a template for exploring the structural topology of states for which no atomic structure is currently available: namely, the MgADP-bound state.

In order to directly correlate spectroscopically-derived results with protein structure, our first consideration was the choice of initial reference model for actomyosin. A recent cryo-EM model of human cytoplasmic myosin-IIc bound to actin was chosen for use as a template (PDB ID: 5JLH), due to the extremely high resolution of the map compared to previous models using myosin-II, and the excellent structural alignment of the derived coordinates with those previous models (Fig. S3A) (34–36). In order to examine this topology in the context of our *Dictyostelium* myosin constructs, we generated a homology model of the *Dictyostelium* sequence using the 5JLH myosin structure as a template (Fig. S3B) (37–40). This homology model was subsequently combined with the F-actin structure from 5JLH by successive structural alignments, yielding a final model consisting of two myosin CD’s bound to adjacent monomers in the same actin polymer chain. In this way, we obtained an actomyosin model capable of representing both distances within an actin-bound myosin CD, and distances between neighboring CD’s on an actin filament.

The second major consideration was the choice of initial starting model for BSL. Only one crystal structure of BSL attached to positions *i* and *i*+4 on an α-helix has currently been reported, using T4 lysozyme as the labeling target (PDB ID: 3L2X) (15). Thus, while the underlying tertiary structure we used for the nucleotide-free actomyosin complex has been supported by near-identical alignments across several isoforms in independent studies, the conformation of BSL reported by crystallography is less substantiated (8, 34, 35, 41). Our previous work has strongly suggested that BSL tends to adopt a single primary conformation on straight α-helices, but that this conformation is not truly represented in the lysozyme crystal structure, due to the placement of labeling sites near a kink in the secondary structure (16). Therefore, the first question we sought to answer with our model concerns optimization of spin label conformation. Specifically, we asked whether there exists a single, consistent conformation of BSL which, when modeled onto each helix in our actomyosin structure, is sufficient to generate agreement with the primary distance and orientational distributions gathered thus far from EPR.

We began by modeling BSL onto each of the three experimentally-labeled helices at appropriate positions in both myosin CDs, by least-squares alignment of backbone atoms from the published coordinates in 3L2X (15). Additional minimization was performed after this initial alignment to ensure that each label had precisely the same spatial orientation relative to its associated myosin helix (see Supplemental Information).

Next, eight values from the nucleotide-free EPR experiments were imposed as constraints in a Monte Carlo minimization of BSL’s spatial conformation: three sets of *θ_NA_* and *ϕ_NA_*, the polar angles of the actin filament axis in the nitroxide frame for each labeling site, measured previously (16); *d_K-W_*, the 4.80nm distance center derived from DEER on the HK-to-HW construct; and *d_relay-W_*, 2.64nm distance center derived from DEER on the relay-to-HW construct (Table 2). During each step of the minimization, the coordinates of all six labels in our model were simultaneously subjected to a geometric transformation *T*, which takes a set of coordinates {*c_n_*} corresponding to the atoms of BSL. The transformation applies an arbitrary rotation Ω*_N_*(*α_N_, β_N_, γ_N_*) about the nitroxide nitrogen, an arbitrary rotation Ω*_R_*(*ψ*) about the helix axis, and an arbitrary translation Ω*_T_*(*a, b*) relative to the helix core and along its axis, thus rendering all possible spatial adjustments to label conformation relative to each helix (*α_N_, β_N_, γ_N_* are Euler angles, *ψ* is an angle about the helix axis relative to the label attachment points, and *a* and *b* are vectors).

**Table 2.** Comparison of experimental and model-derived metrics for the apo actomyosin complex.

| Parameter | Experimental Value<br>(from EPR) | Pre-Minimized<br>Model Value | Post-Minimized<br>Model Value | Adjustment | Post-Minimized<br>Residual |
| --- | --- | --- | --- | --- | --- |
| $d_{K-W}$ | 48.0 Å | 50.2 Å | 47.7 Å | -2.5 Å | -0.3 Å |
| $d_{Relay-W}$ | 26.4 Å | 26.4 Å | 26.2 Å | -0.3 Å | -0.2 Å |
| $\theta_K$ | 88.2° | 93.9° | 85.1° | -8.9° | -3.1° |
| $\phi_K$ | 33.3° | 50.2° | 34.9° | -15.3° | +1.7° |
| $\theta_{Relay}$ | 109.5° | 105.3° | 111.5° | +6.3° | +2.0° |
| $\phi_{Relay}$ | -26.9° | -18.3° | -29.4° | -11.1° | -2.5° |
| $\theta_W$ | 152.0° | 155.7° | 152.5° | -3.2° | +0.5° |
| $\phi_W$ | 147.4° | -168.4° | 148.2° | +43.4° | +0.8° |

Experimental orientation values reported here are equivalent to the mean orientational distribution values given previously (16), with symmetric reflections applied to some values to account for the intrinsic degeneracy in EPR's sensitivity to $\theta_{NA}$ (twofold) and $\phi_{NA}$ (fourfold).

Least-squares optimization of the model against all eight experimental constraints thus corresponds to minimizing the function

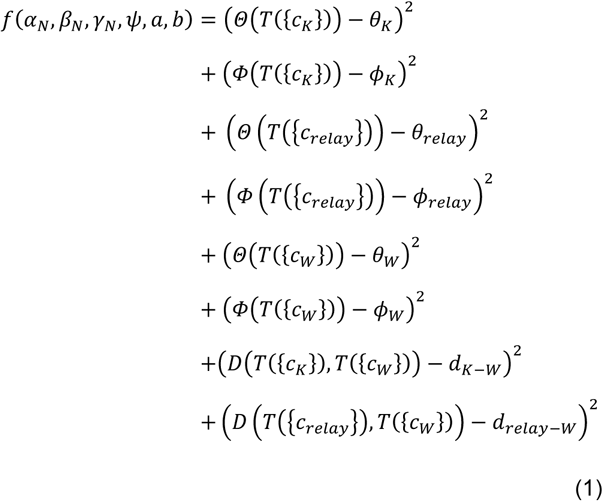

where {*c_n_*} is a set of atomic coordinates corresponding to a spin label at location *n*; *θ_n_* and *ϕ_n_* are experimental *θ_NA_* and *ϕ_NA_* values for location *n*; and *Θ, Φ*, and *D* are functions for calculating *θ_NA_*, *ϕ_NA_* and interspin distances, respectively, from the model. A minimum was found with the following parameters: *α_N_* = −91.3°, *β_N_* = −12.9°, *γ_N_* = 105.8°, *ψ* = −2.9°, *a* = −0.3Å, and *b* = −2.9Å; the optimal transformation *T* was subsequently applied to the coordinates of each BSL molecule.

A comparison between experimental constraints and equivalent measurements in our optimized model is given in Table 2, and the pre-and post-optimization label conformations are overlaid in Fig. 4. It is striking to observe that a relatively small adjustment to the crystal conformation is sufficient to bring all measured parameters to within 3° and 0.3 Å of experimentally-determined values. In addition to the values reported in Table 2, we also surveyed interspin distances between neighboring myosin molecules along the actin filament, and determined that all possible interactions are in the very long distance regime, beyond 5.5nm; this result demonstrates that our derived model solution does not give rise to any conflicting distances that are absent in our experimental analysis. Thus, a single, consistent label conformation employed within our cryo-EM-based actomyosin model is indeed sufficient to reproduce all primary, experimentally-determined parameters with high precision (Table 2, Fig. S4).

**Fig. 4.**
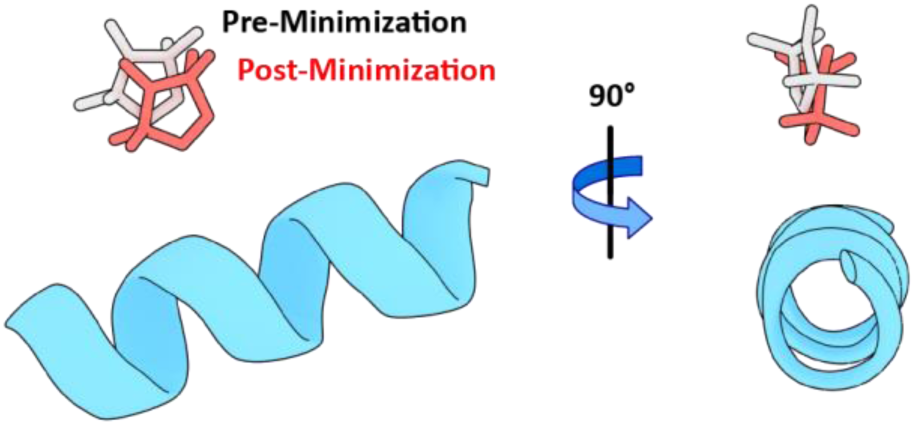
Optimization of spin labels and protein structure in the apo state. (A) Conformation of BSL applied in our actomyosin model, before (white) and after (red) minimization against experimentally-derived constraints.

Next, we turned to the minor sub-populations left unaccounted for in our single-conformation hypothesis, comprising the minor 3.95nm distance observed with the HK-to-HW construct, the minor 3.88nm distance observed in the Relay-to-HW construct, and a pair of *θ_NA_* and *ϕ_NA_* values from a secondary sub-population in the oriented fiber analysis for Helix W reported previously (16). Because Helix W was the only labeling location with two distinct, well-ordered spectral components in oriented fibers, we expanded our analysis to allow for two label conformations on Helix W only. To determine the most likely starting point for optimization of the alternate conformer, we re-examined the crystal structure of BSL on T4 lysozyme used in the previous hypothesis (15). The published coordinates for this structure report a single conformation of the spin label, but examination of the associated electron density map reveals significant patches of unfitted density that likely correspond to a minor secondary conformation (Fig. S5). A coarse atomic model of this minor conformation was generated by rigid-body fitting of BSL coordinates into the unfitted density from 3L2X (Fig. S5) (42), and subsequently modeled onto Helix W in our optimized model by the same least-squares alignment described above. A variation on the algorithm in Equation 1 was performed, this time allowing only the alternate conformation to move while the labels from the optimized model at all other locations remained fixed. For experimentally-derived constants *θ_n_*, *ϕ_n_* and *d_n_* values corresponding to secondary sub-populations were supplied in place of their primary counterparts. Once again, agreement with all constraints was found with minimal adjustment of the starting conformation (Fig. S4, Table S1). Combined with the analysis detailed above, this result serves to account for all populations observed in EPR.

Thus, the cryo-EM-based model we used to represent the nucleotide-free rigor complex is validated by our experimental and modeling results. Furthermore, our hypothesis regarding the highly stereoselective nature of BSL labeling is strongly supported by agreement between our labeled model and experimental results obtained using two different EPR techniques across multiple labeling sites.

### The structure of the MgADP-bound rigor complex determined by application of experimental constraints

Having validated our structure of spin-labeled actomyosin in the apo state, we applied additional orientation and distance constraints to determine how this structure changes in the presence of saturating MgADP. Our approach was similar to the BSL conformer optimization method described above; this time, however, adjustments to labeled helices were allowed while spin label conformation was held fixed.

Several studies have provided strong evidence for a structural transition coupled to ADP release in the actomyosin rigor complex, and myosin-only crystal structures also report subtle differences in protein conformation (see Discussion). However, the sensitivity of these techniques has been largely limited to changes in the force-generating region of S1: the only significant differences observed in crystals occur at the interface between the L50 and Converter domains, and while force-sensing and motility studies reveal lever arm movement in several myosin isoforms, changes internal to the CD cannot be directly observed by these methods. Thus, our constrained modeling approach was designed to allow all three labeled helices full freedom of movement, comprising 3 translational modes and 3 rotational modes. Helices K and W are straight α-helices in all reported crystal structures, with negligible movement relative to surrounding tertiary structure across various biochemical conditions; therefore, these helices were assumed to behave like rigid rods, with rotational pivot points placed at their centers of geometry. The relay helix, on the other hand, has been observed in crystals to exhibit a kink at residue 486, which is strongly coupled to bound nucleotide; thus, only the C-terminal region of the helix (comprising residues 486-496) was allowed to move, with a rotational pivot centered at residue 486.

Once again, eight constraints were applied in our structural refinement: namely, the same set of six angles and two distances employed in the previous section. This time, however, the values of these constraints were derived from experiments performed in the presence of 5mM MgADP (Table 1, Table 3), and the starting model for refinement was the final, optimized model of the apo state derived in the previous section. Choice of this model for the initial condition necessarily biases our refinement toward the solution which most closely resembles the apo complex, but this inherent assumption is justified by the subtlety of the internal changes expected. During optimization, all atoms in each helical segment (including attached spin labels) were subjected to the transformation:

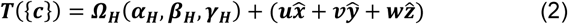

where {***c***} is a set of atomic coordinates, {***u, v, w***} are scalar coefficients; 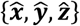 are unit vectors corresponding to principle axes of the global coordinate system; and Ω_H_ defines a rotation described by *α_H_, β_H_*, and *γ_H_*, which are the yaw, pitch, and roll angles associated with each helix in the global coordinate frame, respectively. The optimal transformation was found by least-squares minimization of Equation 1, using Equation 2’s definition of *T* and experimentally-derived constraints corresponding to MgADP experiments, rather than those in the absence of nucleotide. Unlike the spin label optimization in the previous section, which enforced equivalent transformations across all labeling sites, here each helical segment was allowed to move independently, resulting in a unique set of 6 transformation parameters for each of the three helices considered.

**Table 3.** Comparison of experimental and model-derived metrics for the MgADP-bound actomyosin complex.

| Parameter | Experimental Value<br>(from EPR) | Pre-Minimized<br>Model Value | Post-Minimized<br>Model Value | Adjustment | Post-Minimized<br>Residual |
| --- | --- | --- | --- | --- | --- |
| $d_{K-W}$ | 48.0 Å | 47.7 Å | 48.1 Å | +0.4 Å | +0.1 Å |
| $d_{Relay-W}$ | 26.4 Å | 26.2 Å | 26.4 Å | +0.2 Å | 0.0 Å |
| $\theta_K$ | 89.1° | 85.1° | 89.9° | +4.8° | +0.7° |
| $\phi_K$ | 35.2° | 34.9° | 36.8° | +1.9° | +1.6° |
| $\theta_{Relay}$ | 119.1° | 111.5° | 119.1° | +7.6° | 0.0° |
| $\phi_{Relay}$ | -13.6° | -29.4° | -13.3° | +16.1° | +0.2° |
| $\theta_W$ | 155.0° | 152.5° | 154.9° | +2.5° | 0.0° |
| $\phi_W$ | 157.4° | 148.2° | 157.7° | +9.4° | +0.3° |

Optimal parameters from the minimization are given in Table S2. Pre-and post-MgADP-minimization models are overlaid in Fig. 5A-C, and a comparison of experimental measurements to those calculated from the post-minimization model is given in Table 3. Strikingly, these results show that without influence from any existing structures of the MgADP state, the relay helix in our model converges to a conformation which agrees remarkably well with myosin-only crystal structures (Fig. 5D). Results from our previous study also support this notion of structural similarity between actin-free and actin-bound states of the relay helix (16), but here the additional distance constraints from DEER make it possible to arrive at this conclusion completely independently. Furthermore, it is now possible to precisely probe additional changes that were previously observed with orientation measurements alone: Helix W is also altered by the presence of MgADP (Fig. 5D). While Helix K remains almost entirely static, Helix W experiences a very slight shift relative to the center of the actin-binding cleft, with residues closer to the actomyosin interface angling away (Fig. 5A-B). If this observation is assumed to represent bulk movement of the Lower 50kDa domain, it would represent a subtle opening of the actin-binding cleft upon MgADP binding. To our knowledge, this is the first quantified report of a MgADP-induced effect on the structural state of the cleft.

**Fig. 5.**
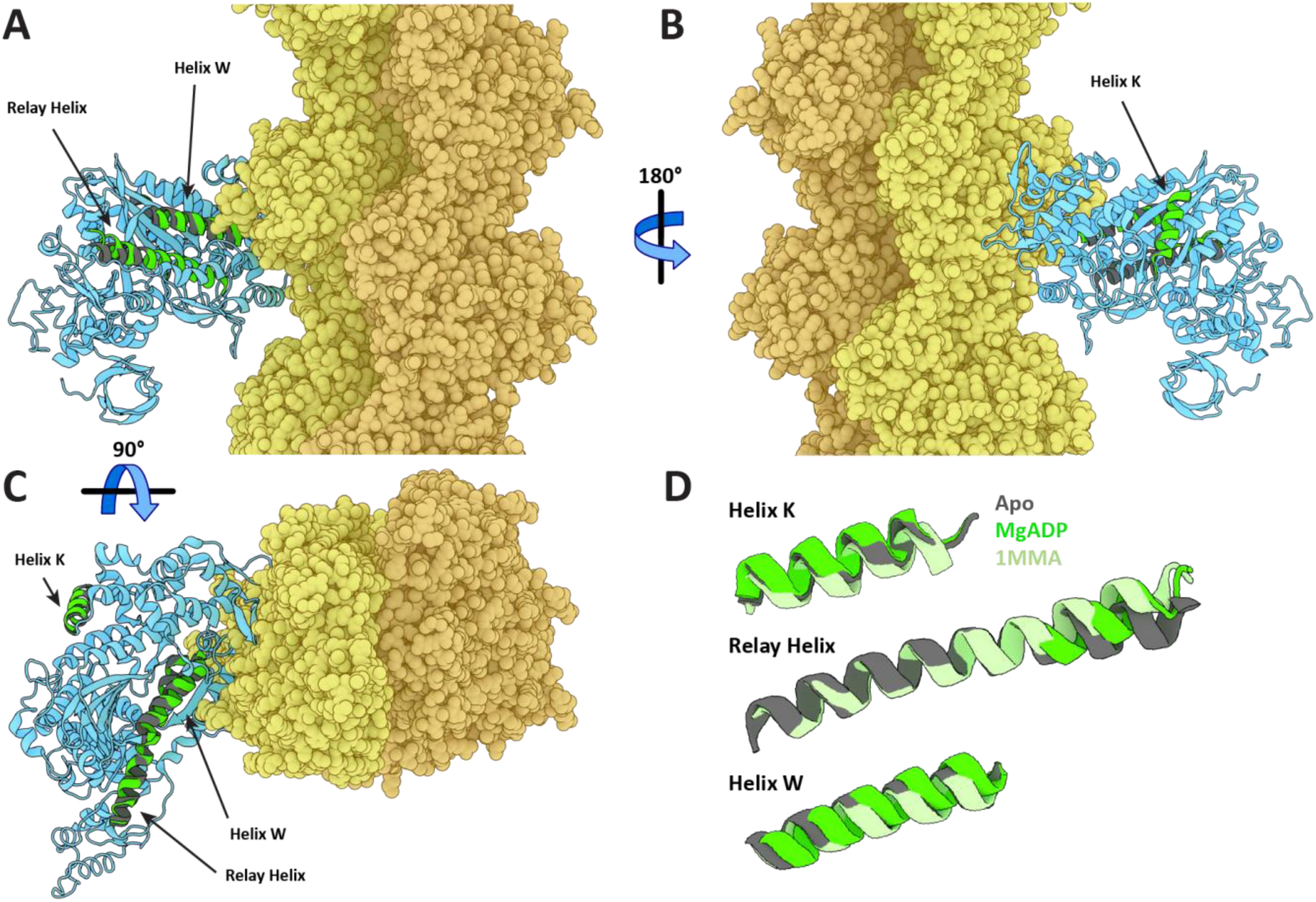
Refinement of the MgADP-bound state of actomyosin. A-C: Myosin (blue) and actin (yellow/orange) from our model of the rigor complex, before and after MgADP refinement. Targeted helices (K, relay, and W) are highlighted in the pre-refinement (apo state, dark gray) and post-refinement (MgADP state, green) states. D: Comparison of the apo state (dark gray) and MgADP state (green) models to a crystal structure of Dictyostelium myosin without actin, bound to MgADP (PDB ID: 1MMA, light green) (43). Because of significant differences in the overall arrangement of domains around the actin-binding cleft between crystal (cleft open) and model (cleft closed) structures, alignment of the crystal structure to the model was accomplished via pair-wise matching of either the Upper 50kDa domain (for Helix K) or the Lower 50kDa domain (for the Relay Helix and Helix W).

## Discussion

### BSL adopts a highly stereoselective conformation on protein α-helices

BSL has been employed as an attractive alternative to monofunctional spin labels in a variety of systems, but while several attempts have been made to characterize the label’s binding behavior, definitive conclusions are difficult to draw (15, 17, 18, 44). Pure *in silico* analysis tends to produce conformational distributions which are significantly wider than those observed in experiments (16, 18, 44), and the only crystal structure currently available is unlikely to reflect a preferred conformation on straight α-helices due to the selected positioning of reactive cysteines in the target protein (15). Thus, it has been very challenging to relate BSL-derived measurements of orientation and distance directly to the behavior of an associated protein backbone, despite much evidence that the label readily becomes strongly immobilized and highly ordered upon bifunctional attachment.

In the present study, we have demonstrated that the advantages of BSL for increased distance resolution in DEER still apply when employed in the large and dynamic myosin CD (Fig. 1). By examining multiple spin label pairs across the CD, we have determined that BSL produces distance distributions of a width comparable to the width observed in orientational distributions from oriented fibers (Fig. 2, Fig. S1, Fig. S2, Fig. S4, Table 1) (16).

Crucially, we have used this experimental data to determine a single label conformer that is consistently preferred across three separate labeling sites on different helices. Thorough *in silico* modeling and optimization reveals that a single conformation is sufficient to satisfy all orientation and distance constraints from our EPR experiments (Fig. 4, Fig. S4) (16, 36). This label conformation is significantly different from the published crystal coordinates, though in comparison, the label still retains its position on the same side of the helix, and its orientation relative to the helix axis is similar (15). We have also characterized a secondary conformation, which is sufficient to account for all significant minor sub-populations in experimental distributions (Fig. S4). Results from our optimization indicate that this alternate conformation is likely present on just one of the three helices, and that it is similar to the conformation suggested by unfitted density in the published crystallographic data (Fig. S5, Table S1) (15).

Tight coupling of spin labels to the protein backbone was a goal that inspired the original synthesis of BSL (13), and this property has since been well-established in the literature (5, 15, 17, 44, 45). In the present work, we expand upon a hypothesis from our previous study, demonstrating that BSL is not only strongly immobilized and well-ordered on α-helices, but also *highly stereoselective.* Our results indicate that the label takes at most two conformations on the three helices surveyed, and that one of these conformations is strongly preferred over the other. The notion of stereospecificity in site-directed spin labeling represents tremendous potential for high-resolution structure determination, because the relationship between backbone and nitroxide is unambiguous, and therefore subject to direct modeling. We have determined structures for each of the observed conformations, and thus our findings are readily applicable to other protein systems via direct modeling.

### The C-terminal end of myosin’s relay helix rotates upon binding of MgADP when in complex with actin

The ADP-release step in myosin’s catalytic cycle has garnered increasing attention as a source of important coupled structural changes in Subfragment-1, particularly as they pertain to force generation. Movement of the myosin lever arm at a rate comparable to the release of ADP has been observed in several isoforms (46–49), as have strain-dependent effects on ADP release rate (48–54). Crystal structures of myosin alone have long predicted a movement of the C-terminal end of the relay helix upon binding of various nucleotides, including MgADP, which are thought to correlate with large-scale transitions of the lever arm (28–30, 43, 55). These interactions have been difficult to observe in actin-bound myosin, but examples for processive nonmuscle and smooth muscle myosins are available (33, 56).

We have demonstrated previously that a structural transition also occurs in actin-bound myosin-II upon MgADP binding/release: results from EPR on oriented fibers indicated movement of the C-terminal end of the relay helix that was likely to mirror the shift observed in myosin-only crystal structures (16). However, our conclusions were limited by the relatively small number of experimental constraints, such that we were only able to quantitatively report the change in helix tilt angle relative to the actin filament axis.

In the present study, our experimental constraints for this system have been significantly expanded with the addition of DEER-derived distance distributions. The additional parameters have allowed us to render a complete geometric transformation for the observed changes in the relay helix, resulting in a new set of coordinates that may be compared to myosin-only crystal structures (Fig. 5). Our final model falls in excellent agreement with these structures, indicating that the presence of actin does not significantly alter the structural mechanism of force propagation in the myosin CD.

### Changes in the actin-binding cleft upon binding of MgADP

Many studies have shown that the changes to myosin induced by MgADP are not limited solely to the force-generating domain. The binding of MgADP in myosin’s nucleotide binding pocket has been shown to drastically reduce the affinity of myosin for actin, especially for classes characterized by a low duty cycle, like myosin-II (57–61). This change in affinity implies that structural transitions also occur near the actomyosin interface as a result of ADP binding/release, but no study has yet determined the extent or nature of these changes in myosin-II.

The actin-binding cleft is a large solvent-filled cavity between the Upper and Lower 50kDa domains in myosin’s CD that has been widely observed to narrow (or “close”) when myosin transitions from a weak-binding interaction into a strongly-bound rigor complex with actin (8, 33–36, 41, 62, 63); this correlation has led to the adoption of cleft conformation as a reporter for actomyosin binding affinity in structural studies (64). We have previously demonstrated that Helix W, located near the cleft in the Lower 50kDa domain, experiences a significant shift in orientation relative to actin upon binding of MgADP by myosin (16). However, just as with the relay helix, analysis of this change was limited by a lack of sufficient modeling constraints.

In the present study, additional constraints from DEER have allowed for a more complete modeling of changes to both Helix K and Helix W with MgADP binding/release. Both helices remain very close to their original positions, but Helix W experiences a subtle shifting of its axis relative to actin, such that the residues closest to the actin interface are angled slightly away. Residues on Helix W are expected to form direct electrostatic contacts with actin, and this shift alone would cause potentially significant changes to that interface. Investigation of additional sites on the Lower 50kDa domain would further elucidate these changes.

It is worth noting that the reciprocal actin/MgADP affinity effect noted above is blunted by at least an order of magnitude in the *Dictyostelium* construct that serves as our background for site-directed labeling. In wild-type *Dictyostelium* myosin, MgADP affinity is reduced roughly 100-fold upon actin binding, but substitution of native cysteines and truncation of the construct at residue 758 are both correlated with drastic attenuation of this effect (61, 65–67). Thus, it is possible that the effect we observe here is less pronounced than it would be in a wild-type motor. In the future, additional studies using longer myosin constructs would be invaluable for investigating the apparent allosteric coupling mediated by the C-terminus; optimization of labeling methods may also permit the study of constructs with some native cysteines restored.

## Conclusions

In this study, we combined two complementary EPR methods using a bifunctional spin label to model the structure of myosin-II when bound to actin filaments. We determined that the use of BSL for the study of myosin with DEER provided superior distance resolution compared to monofunctional labels (Fig. 1), and measured distances across the myosin CD (Fig. 2 and Fig. 3). Molecular modeling showed that a single conformer of BSL was sufficient to model all distances and orientations derived from EPR experiments (Fig. 4) (16), strongly supporting our conclusion that BSL is highly stereoselective on protein α-helices. Minimization against orientation and distance constraints derived from experiments, including MgADP allowed for direct modeling of specific changes to individual structural elements within the CD when MgADP is bound (Fig. 5). The Relay Helix exhibits changes that mirror its behavior in myosin-only crystals, while Helix W exhibits subtle movements that were not possible to model directly with orientation measurements alone. Collectively, this work demonstrates the power of site-directed spectroscopy using stereospecific spin labels, and outlines a method for applying EPR-derived orientation and distance constraints directly to atomic models.

## Author Contributions

B.P.B and D.D.T. designed the research; B.P.B. carried out the majority of the spectroscopy experiments; A.R.T. performed additional experiments; B.P.B., A.R.T., and D.D.T. analyzed resulting waveforms; B.P.B. performed the molecular modeling analysis; B.P.B. and D.D.T. wrote the manuscript. All authors approved the final version of the manuscript.

## Acknowledgements

EPR experiments were performed at the Biophysical Technology Center, University of Minnesota. This study was supported by National Institutes of Health (NIH) Grants AR32961 and AG26160 (to D.D.T.). B.P.B. was supported by NIH Grants T32 AR007612 and K12 GM119955.

